# Transcriptomic analysis of astaxanthin hyper-producing *Coelastrum* sp. mutant obtained by chemical mutagenesis

**DOI:** 10.1101/2021.08.17.456660

**Authors:** Ameerah Tharek, Shaza Eva Mohamad, Iwane Suzuki, Koji Iwamoto, Hirofumi Hara, Shinji Yoshizaki, Haryati Jamaluddin, Madihah Md Salleh, Adibah Yahya

**Author notes:** Corresponding author Associate Professor Dr. Shaza Eva Mohamad, Malaysia Japan International Institute of Technology (MJIIT), Department of Chemical and Environmental Engineering (CHEE), Universiti Teknologi Malaysia, Jalan Sultan Yahya Petra, 54100, Kuala Lumpur, Malaysia.

## Abstract

A newly isolated green microalga, *Coelastrum* sp. has the capability to produce and accumulate astaxanthin under various stress conditions. At present, a mutant G1-C1 of *Coelastrum* sp. obtained through chemical mutagenesis using ethyl methane sulfonate displayed an improvement in astaxanthin accumulation, which was 2-fold higher than that of the wild-type. However, lack of genomic information limits the understanding of the molecular mechanism that leads to a high level of astaxanthin in the mutant G1-C1. In this study, transcriptome sequencing was performed to compare the transcriptome of astaxanthin hyper-producing mutant G1-C1 and wild-type of *Coelastrum* sp. with respect to astaxanthin biosynthesis. This is to clarify why the mutant produced higher astaxanthin yield compared to the wild-type strain. Based on the transcriptomic analysis, the differentially expressed genes involved in astaxanthin biosynthesis were significantly upregulated in the mutant G1-C1 of *Coelastrum* sp. Genes coding phytoene synthase, phytoene desaturase, *ζ*-carotene desaturase, and lycopene β-cyclase involved in β-carotene biosynthesis in the mutant cells were upregulated by 10-, 9.2-, 8.4-, and 8.7-fold, respectively. Genes coding beta-carotene ketolase and beta-carotene 3-hydroxylase involved in converting β-carotene into astaxanthin were upregulated by 7.8- and 8.0-fold, respectively. In contrast, the lycopene ε-cyclase gene was downregulated by 9.7-fold in mutant G1-C1. Together, these results contribute to higher astaxanthin accumulation in mutant G1-C1. Overall, the data in this study provided molecular insight for a better understanding of the differences in astaxanthin biosynthesis between the wild-type and mutant G1-C1 strains.

## Introduction

Astaxanthin is an increasingly popular antioxidant that belongs to a family of naturally occurring pigments called carotenoids. It is regarded as a powerful natural antioxidant compared to other carotenoids [1]. Nowadays, the interest in astaxanthin is rising due to its superior antioxidative activity that benefits human health. Due to the potential of health benefits, its demand is highly increased in nutraceutical, pharmaceutical, cosmetics, and aquaculture industries [2–5].

Thus, significant efforts have been made in recent years to improve astaxanthin production to meet the rising market demand. Although astaxanthin is ubiquitous in nature, its biosynthesis is limited to some microorganisms. Astaxanthin biosynthesis has been observed to be commercially cultivated in a limited number of organisms such as bacteria, yeast, fungi, higher plants, and some green microalgae [6, 7]. Microalgae are currently known to have the highest level of astaxanthin accumulation in nature by being the primary source of natural astaxanthin for human consumption. However, the market supply of natural astaxanthin is still unable to satisfy global demand [3].

Therefore, there have been numerous efforts to improve microalgae strains with high astaxanthin yields. Multiple routes have been explored to produce astaxanthin by selecting high-yield strains, optimizing the cultivation, using chemicals as metabolic enhancers, and genetically modifying the strains [8–11]. Among the various methods in improving astaxanthin, the random mutagenesis technique has been identified to effectively generate high astaxanthin accumulation in microalgae [12].

Random mutagenesis technique has been widely applied to improve the productivity of various algal strains by constructing genetically improved mutants with excellent growth rate and high astaxanthin production [13–16]. A study by Sandesh Kamath et al. (2008) showed improvement of total carotenoid and astaxanthin contents by 23–59% in *Haematococcus* mutant exposed to mutagens of ultraviolet light (UV), ethyl methane sulphonate (EMS), and 1-methyl 3-nitro 1-nitrosoguanidine (NTG) [13]. Chen et al. (2003) also isolated *H. pluvialis* mutant by UV and EMS, and it accumulated higher astaxanthin production in the mutant [17]. Besides that, a genetic improvement to a Chilean strain of *H. pluvialis* mutated by EMS was found to improve its carotenogenic capacity by 30% more astaxanthin compared to the wild type strain [18].

Presently, the discovery of astaxanthin in *Coelastrum* sp. strain has attracted attention due to its potential in accumulating astaxanthin [8, 19, 20]. To date, the genetically modified *Coelastrum* sp. strain has not been reported for commercial production of astaxanthin. In this study, a newly isolated strain of *Coelastrum* sp. demonstrated enhanced astaxanthin accumulation was produced via genetically modified method by chemical mutagenesis using EMS. However, the genetic information and the molecular mechanism of astaxanthin synthesis routes resulting in high astaxanthin production in *Coelastrum* sp. mutant are still unknown. Therefore, to improve our understanding of astaxanthin biosynthesis in *Coelastrum* sp., a comprehensive analysis using RNA sequencing (RNA-seq) have been employed. Transcriptome sequencing of microalgae culture with differential gene expression analysis would be an efficient approach to explore the mechanism of astaxanthin synthesis in *Coelastrum* sp.

RNA-seq is a recent application of the sequencing technique for analyzing the whole transcriptome structure and transcription level of individual genes [21, 22]. This approach will greatly improve our understanding of astaxanthin biosynthesis in *Coelastrum* sp. In the present study, the comparative transcriptomic analysis of the gene expression profile was conducted by comparing the wild-type and astaxanthin hyper-producing mutant to examine differential regulations of genes associated with particular pathways of interest. The data from transcriptomic analysis can be used to identify all genes involved in astaxanthin biosynthesis of *Coelastrum* sp. This will act as a vital resource for improving astaxanthin production and provide a new insight to expand our understanding of *Coelastrum* sp. toward carotenoid metabolism and biosynthesis.

## Materials and methods

### Strains and culture conditions

The wild-type of newly isolated *Coelastrum* sp. isolated from a sampling site at Hulu Langat river, Kuala Selangor, Malaysia, and *Coelastrum* sp. mutant G1-C1 strain obtained through chemical mutagenesis approach (0.4 M of EMS with 60 min exposure time) were cultured in AF-6 medium according to media recipe available in the Microbial Culture Collection National Institute for Environmental Studies (NIES-collection), Japan [23]. The cell cultures were grown under controlled laboratory conditions at 25±1 °C, illuminated at a continuous light intensity of 70 μmol photons m^−2^ s^−1^, and aerated continuously with 1% CO_2_ until it reached the exponential growth phase.

The green cells of wild-type and mutant strain were then harvested. Various supplements were added according to optimize conditions in accumulating astaxanthin in *Coelastrum* sp. with details described in our previous work [10]. Sodium acetate, sodium chloride, and sodium nitrate were used at a final concentration of 0.5 g/L, 3 g/L, and 0.1 g/L, respectively, and were subsequently exposed under continuous illumination of high photon flux densities of 250 μmol photon m^−2^ s^−1^ for the induction of astaxanthin biosynthesis. The cells were then collected through centrifugation and stored at −80 °C until further analysis. All the experiments were carried out in triplicates.

### Measurement of dry cell weight

Dry cell weight was measured based on Boussiba and Vonshak (1991) method. The empty centrifuge tube was placed in an oven for 30 min to remove excess moisture and weighed. Then, 5 mL of culture fluid was centrifuged at 5000 ×*g* for 10 min at 4 °C. The pellet was re-suspended in distilled water, dried at 70 °C in the oven until a constant weight was obtained, and cooled down to room temperature before weighing. The biomass weight was expressed in g/L [24].

### Extraction and analysis of total carotenoid and astaxanthin

To measure total carotenoid and astaxanthin content, 15 mL volume of wild type and mutant culture was centrifuged at 2000 ×*g* for 10 minutes at 4 °C. The pellet was lyophilized using a freeze dryer (Lyphlock 6; Labconco, USA). Then, the cells were homogenized with acetone and kept in a water bath at 70 °C for 10 min followed by vortexing for few minutes. The mixture was centrifuged at 2000 ×g for 10 min and the supernatant was collected. Supernatant collections were conducted repeatedly until the cells were faded. The concentration of total carotenoid was estimated by measuring at absorbance 470 nm and calculated using the Lichtenthaler (1987) equations [25]. The astaxanthin concentration was then measured by the spectrophotometric method and calculated with the equation, c (mg/L) = 4.5 × A_480_ × (V_a_ / V_b_) × f. Where c is the astaxanthin concentration, V_a_ (mL) is the volume of solvent, V_b_ (mL) is the volume of algal sample, and f is the dilution ratio. 480 nm was the absorption peak of astaxanthin. A_480_ was determined by measuring the absorbance at 480 nm. Acetone was used as blank for the measurement.

### RNA extraction and sequencing

Total RNA was extracted from the wild-type and mutant G1-C1 cells of *Coelastrum* sp. which were harvested during astaxanthin accumulation induction stages. Approximately 1×10^8^ cells of aliquots were collected by centrifugation, frozen in liquid nitrogen, and subsequently grounded using mortar and pestle into a fine powder and then stored at −80 °C prior to RNA extraction. Trizol reagent was used to extract the total RNA of microalgae cells. The cells were lysed in a 1 mL Trizol reagent and stored in the icebox for 5 min. A total of 200 µL of chloroform was added to the supernatant, and the mixture was shaken vigorously for 15 secs and stored for 10 min in the icebox. Then, the mixture was centrifuged at 12 000 ×g for 15 min at 4 °C, and the upper aqueous phase was carefully transferred into a new 1.5 mL Eppendorf tube. Subsequently, 500 µL isopropanol was added to precipitate the total RNA. The mixture was kept in the icebox for 10 min and centrifuged at 12 000 ×g for 10 min at 4 °C. The supernatant was removed, and the pellet was washed with 1 mL of 75% ethanol. The mixture was then centrifuged (5000 ×*g* for 5 min at 4 °C) and ethanol was removed. The RNA pellet was then briefly air-dried for 5-10 min and dissolved with 50 µL Diethyl Pyrocarbonate (DEPC) treated distilled water and stored at −80 °C for later use. RNA quality and concentration of wild-type and mutant were determined using the Agilent Bioanalyzer 2100 system (Agilent Technologies, CA, USA). Total RNA (~1 μg) for each sample was used for cDNA library construction and *de novo* transcriptomic analysis conducted by Genewiz, Inc. (Hangzhou, China).

### Transcriptome assembly and functional annotation

To obtain high-quality clean reads, the quality assessment of the sequencing data was initially processed by Cut-adapt (V1.9.1). This was to remove technical sequences including adapters, unknown nucleotides, poly-N strands, filtered off low quality reads (Phred score value <20 at the 3′ or 5′ end), and removed sequences shorter than 75 bp. All filtered reads were then evaluated using FastQC (V0.10.1: http://www.bioinformatics.babraham.ac.uk/projects/fastqc/) to confirm the quality of the data. The clean reads from each library were evaluated with the Q20 and Q30 (the percentage of bases with quality scores, Phred value > 20 and > 30, respectively), GC content (percentage of GC in the read), and N-content (number of undetermined bases, N) [26].

Then, the high-quality filtered reads for genome *de novo* assembly were performed using Trinity (V2.2.0) to generate unigene sequences [27]. The assembly result was connected via sequence clustering into long non-redundant sequences, which were called as unigenes. The number and lengths of the resulting contigs are essential for measuring the quality of assembly. Finally, filtered reads were mapped to unigenes using Bowtie2 (2.1.0), which is used for gene expression analysis, and the reads per kilobase of exon model per million (RPKM) mapped fragments were used to represent gene transcription [28].

Afterward, trimmed reads were annotated with protein databases. Functional annotation of the unigenes was performed by Gene Ontology (GO) and Kyoto Encyclopedia of Genes and Genomes (KEGG) enrichment analysis. GO enrichment and KEGG enrichment analysis were conducted according to transcriptome detection results.

### Quantification of gene expression and differential expression analysis

To quantify the gene expression level of the assembled unigenes, gene expression profiles were estimated using RSEM (v1.2.6) which measures in FPKM (Fragment Per Kilo bases per Million reads) [29]. Based on the gene expression, the differentially expressed genes (DEGs) in *Coelastrum* sp. mutant were identified using DESeq2 (V1.6.3) which was based on a model that analyzes RNA-Seq data with a negative binomial distribution [30, 31]. Then, gene differential analysis was performed using EdgeR (V3.4.6). The results from EdgeR analysis were further analyzed to determine genes with significant differential expression. A False Discovery Rate (FDR) method was used to determine the threshold of the *p-*value. Genes with a fold change greater than two and a *p-*value less than 0.05 were considered statistically significant.

### GO and KEGG pathway enrichment analysis of DEGs

GO and KEGG pathway enrichment analysis were used to determine the functions and enriched pathways of the DEGs. The primary biological functions of DEGs can be determined by GO enrichment analysis. The GO functional enrichment analysis shows the GO terms enriched among DEGs against the genomic background. It thus provides information on how the DEGs are related to certain biological functions. GO analysis mapped all the DEGs to the GO database and then calculate the number of differential genes in each term. The GO term with a corrected *p*-value ≤ 0.05 was considered significantly enriched in the DEGs.

DEGs also underwent KEGG pathway enrichment analysis to determine the DEGs involved in the most essential biochemical metabolic pathways and signal transduction pathways using KEGG metabolic pathway database [32]. Pathway enrichment analysis was performed to find the pathways of the DEGs that are significantly enriched against transcriptome background.

## Results and discussion

### Comparison of wild-type and mutant G1-C1 of Coelastrum sp

To investigate changes in pigment content, the wild-type (WT) and mutant G1-C1 strain of *Coelastrum* sp. were analyzed and compared based on their total carotenoid and astaxanthin content. In this study, mutagenesis using chemical mutagen of EMS was attempted to increase the microalgae biomass and carotenoid production in *Coelastrum* sp. **Fig 1** shows that the mutant G1-C1 tends to have higher growth compared to the WT. The biomass of mutant G1-C1 increased significantly with 1.3-fold more elevated than the WT. As shown in **Fig 2**, the astaxanthin content also revealed that mutant G1-C1 acquired 28.32 mg L^−1^ of astaxanthin content which was approximately 2-fold higher astaxanthin compared to the WT strain (14.5 mg L^−1^).

**Fig 1.**
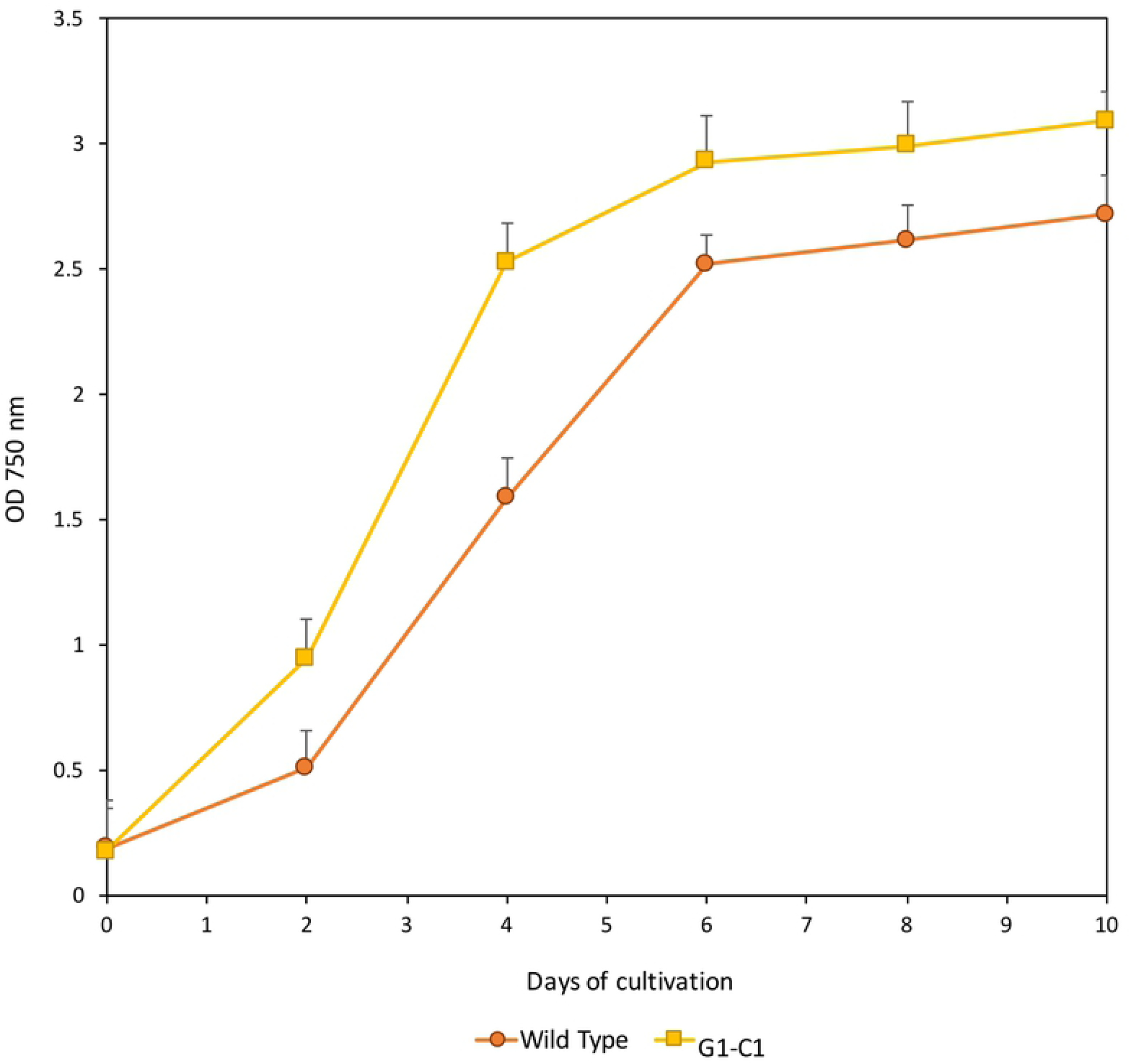
Growth curve of the wild-type and mutant G1-C1 obtained from chemical mutagenesis using chemical mutagen of EMS

**Fig 2.**
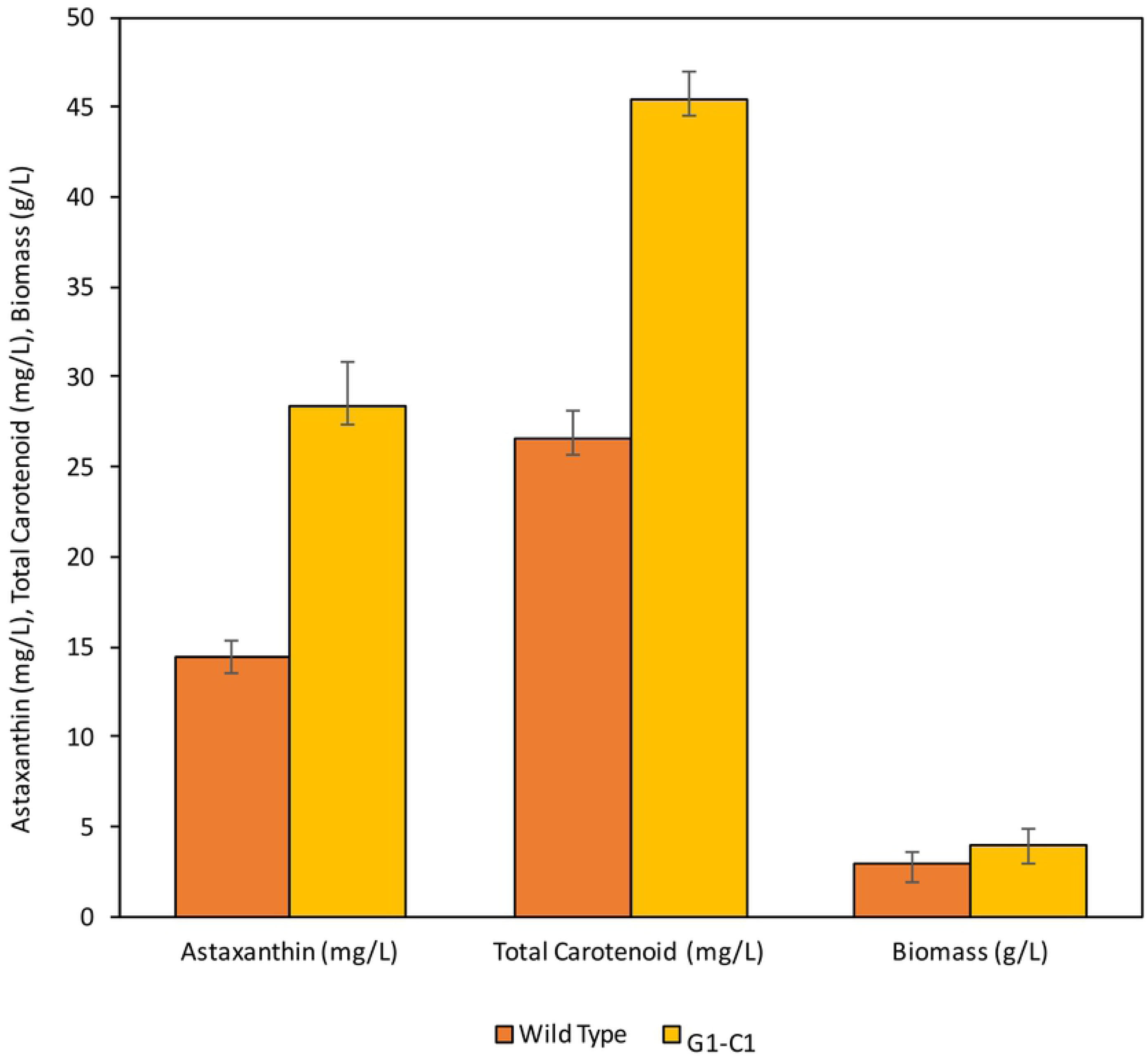
Comparison of biomass, total carotenoids, and astaxanthin content in the wild-type and mutant G1-C1. Data represent an average of 3 replications, and error bars indicate mean ±SD

The findings of this study revealed that the genetic improvement of newly isolated *Coelastrum* sp. by random mutagenesis had altered certain biochemical characteristics of the WT. The altered biochemical properties of *Coelastrum* sp. mutants were demonstrated to be a successful strategy to increase the content of astaxanthin. However, the molecular mechanisms behind these effects are unknown. Therefore, to determine molecular changes and to better understand the high astaxanthin accumulation underlying the mutant G1-C1 strain, a comprehensive analysis using RNA-seq have been employed. The molecular mechanism of the effect of the chemical mutation on astaxanthin accumulation in *Coelastrum* sp. could be determined by studying the changes in gene expression between WT and mutant. A list of genes involved in astaxanthin biosynthesis of *Coelastrum* sp. was also identified using transcriptomic analysis.

### RNA sequencing and de novo transcriptome assembly analysis of Coelastrum sp

Transcriptome sequencing was performed to have a detailed understanding of astaxanthin biosynthesis in *Coelastrum* sp. In the transcriptomic analysis, transcript sequences were obtained from spliced clean reads. The sequencing yielded 85.20 and 88.42 million raw reads for WT and mutant G1-C1, respectively. After removing adapter sequences and low-quality reads, 84.87 million clean reads from the WT, and 85.19 million clean reads from the mutant G1-C1 were obtained. Then, the clean reads were assembled after obtaining high-quality clean reads of data. *De novo* assembly of transcriptomes yield a total of 7,317,366 transcripts that were assembled into 140,551 unigenes with an N50 length of 718 bp. Among the assembled unigenes, 106,151 (75.53 %) were 200-500 bp, 18,874 (13.43 %) were 500-1000 bp, 6,995 (4.98 %) were 1000-1500 bp, 3,572 (2.54 %) were 1500-200 bp, and 4, 959 (3.53%) were more than 2000 bp in length. Summary of RNA sequencing and *de novo* assembly of transcripts presented in **Table 1** and **Fig 3** shows the length distributions of unigenes obtained from *de novo* assembly ranged from 200 to more than 2000 bp. To our knowledge, this study is the first transcriptomic study of *Coelastrum* sp. to be made available. Therefore, the obtained transcript sequences for *Coelastrum* sp. that enrich the genomic resources can be used for gene discovery.

**Fig 3.**
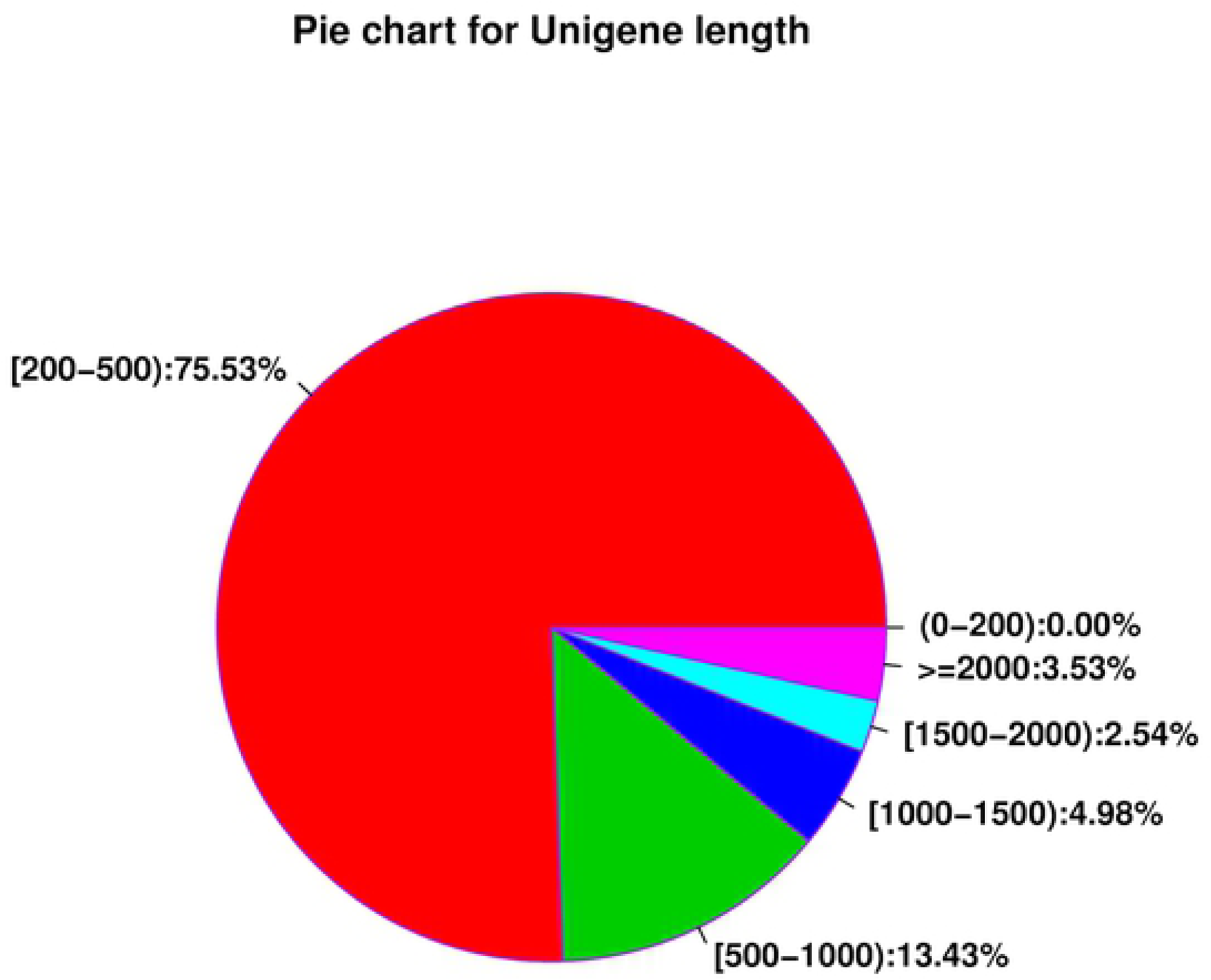
Pie chart of unigenes distribution by length

**Table 1.** Summary of RNA sequencing and *de novo* transcriptome assembly results

|  | Wild Type | Mutant G1-C1 |
| --- | --- | --- |
| Number of raw reads | 85,195,406 | 88,415,780 |
| Average length of raw reads (bp) | 150 | 150 |
| Number of trimmed reads | 84,868,972 | 85,186,382 |
| Number of trinity transcripts | 7,317,366 |  |
| Number of unigenes | 140,551 |  |
| N50 (bp) | 718 |  |
| Number of DEGs with significant expression differences (FDR < 0.05) | 87,892 |  |

### Gene function classification by GO and KEGG

Unigene classification of *Coelastrum* sp. was conducted by GO and KEGG enrichment analysis. A total of 140,551 unigenes were annotated in this study. GO was used to classify the predicted functions of unigenes and standardize the gene functional classification system of each assembled unigenes. GO functional enrichment analysis was divided into three categories, including biological process (BP), cellular component (CC), and molecular function (MF) which included 62 functional subcategories in total in *Coelastrum* sp.

As shown in **Fig 4**, genes of the MF group were divided into 20 subcategories where binding (13.7%) and catalytic activity (13.46%) were the most dominant groups. The genes in the CC category were also divided into 20 subcategories. Cell part (12.79%) represented the majority of this category, followed by membrane part (4.23%), organelle (4.07%), and membrane (4.06%). While, predicted proteins assigned to the BP category were mainly associated with the cellular process (9.66%), metabolic process (9.43%), single-organism process (6.66%), and biological regulation (3.39%). Results have shown that a large fraction of unigenes function differently, and these transcripts could be the genes that control specific cell proliferation and differentiation in *Coelastrum* sp. and thus will be useful for functional gene study [33].

**Fig 4.**
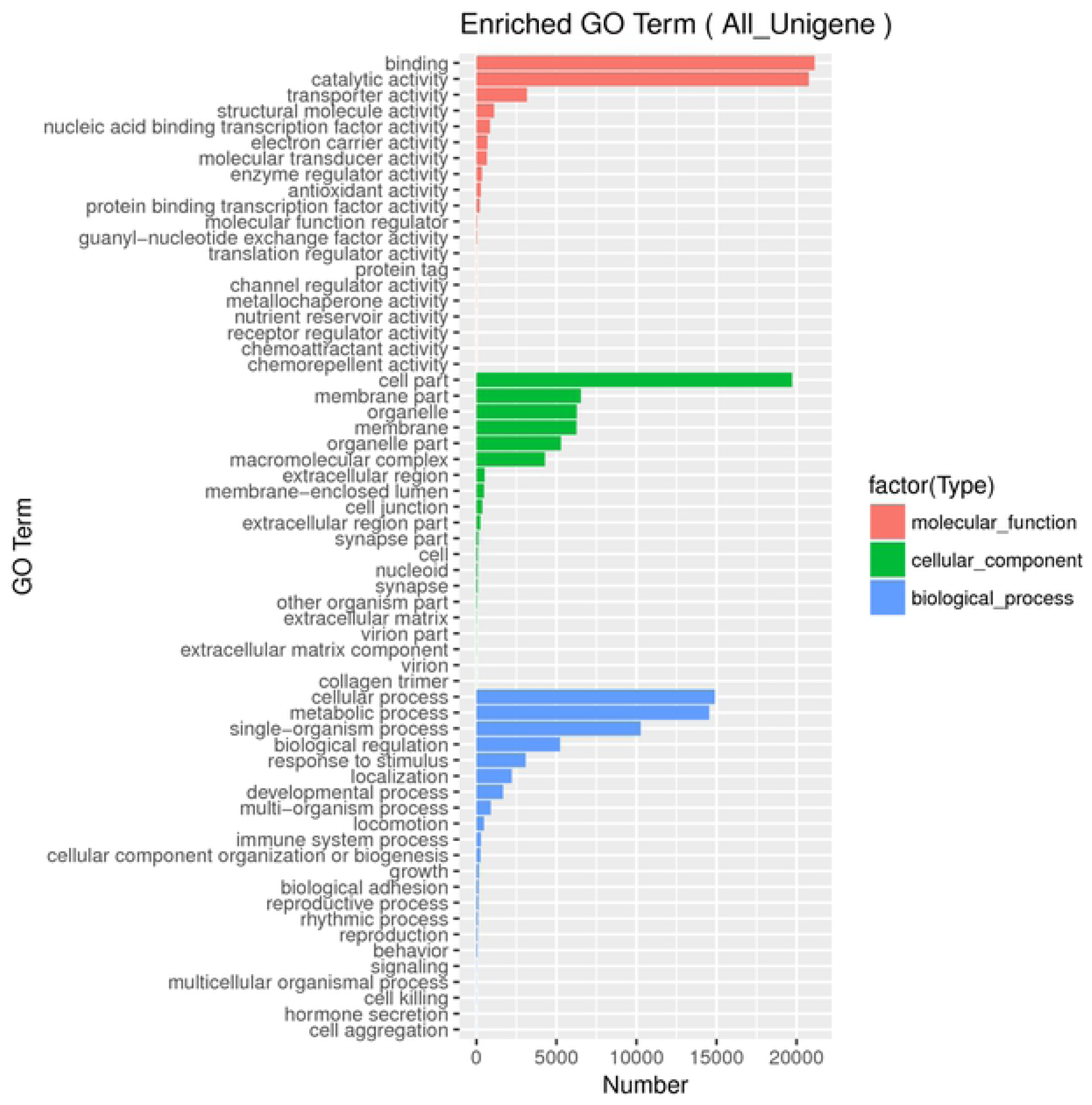
Distribution of annotated sequences based on GO analysis. GO functional classification assigned 140,551 unigenes to 62 subcategories under three main GO categories: molecular function (MF), cellular component (CC), and biological process (BP). Y-axis: the classification of gene ontology; X-axis: the number of genes in each category

Subsequently, after assembling unigenes for functional annotation, all the unigenes were further analyzed via KEGG pathway prediction. KEGG is used for systemic gene functional analysis and genomic studies. It is an approach for categorizing gene functions with an emphasis on biochemical pathways. It integrates genomics, biochemistry, and functionomics to construct biological pathways for different types of biological processes [21].

To interpret the major biochemical pathways and signal transduction involved in *Coelastrum* sp., all the unigenes were analyzed via the prediction of the KEGG pathway. All unigenes were annotated to 20 KEGG sub-pathways (**Fig 5**) which were widely classified in various cellular functions as cellular processes (CP), environmental information processing (EIP), genetic information processing (GIP), human diseases (HD), metabolism (M), and organismal systems (OS). Among these, transcripts associated with metabolism (M) constituted a large fraction in the unigenes indicating a large number of metabolisms involved in *Coelastrum* sp. (**Fig 5**). A total of 14,326 unigenes were involved in global and overview maps which comprised most of the unigenes classified in various metabolic pathways (7,192) followed by biosynthesis of secondary metabolites (3, 356) (**Table 2**). KEGG metabolic pathway analysis showed that the biosynthesis of secondary metabolites is related to astaxanthin biosynthesis. This may contribute to better understanding and further elucidation of the biological functions of astaxanthin biosynthesis in *Coelastrum* sp.

**Fig 5.**
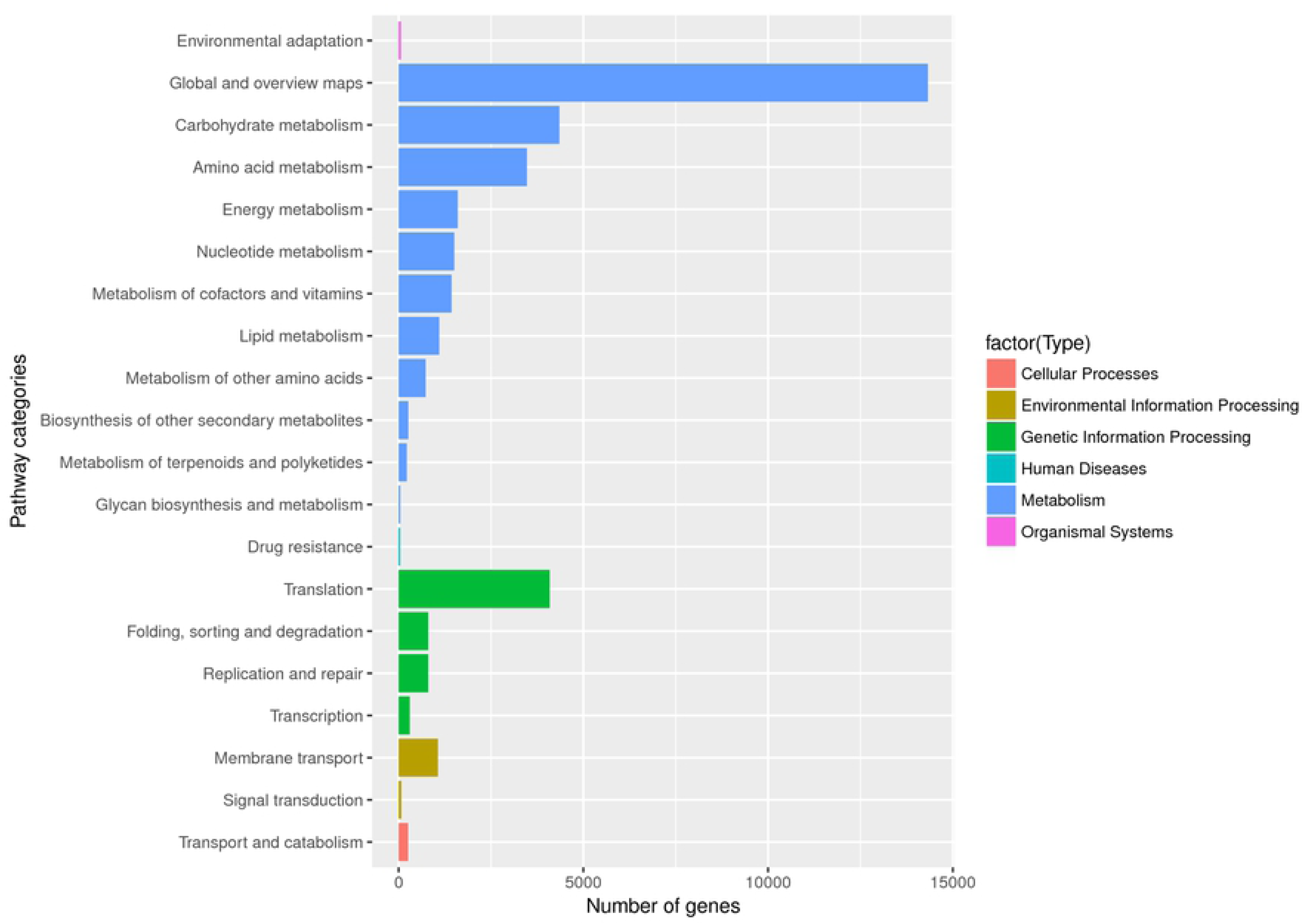
Distribution of annotated sequences based on KEGG pathway analysis. Y-axis: KEGG pathway classification; X-axis: number of genes within each KEGG pathway

**Table 2.** Functional categorization of assembled unigenes in KEGG pathways

| KEGG sub-pathways | Number |
| --- | --- |
| **Metabolism** |  |
| **Global and overview maps** | **14326** |
| KO01100 Metabolic pathways | 7192 |
| KO01110 Biosynthesis of secondary metabolites | 3356 |
| KO01200 Carbon metabolism | 1551 |
| KO01210 2-Oxocarboxylic acid metabolism | 378 |
| KO01212 Fatty acid metabolism | 418 |
| KO01230 Biosynthesis of amino acids | 1431 |
| **Carbohydrate metabolism** | **4351** |
| KO00010 Glycolysis / Gluconeogenesis | 489 |
| KO00020 Citrate cycle (TCA cycle) | 462 |
| KO00030 Pentose phosphate pathway | 249 |
| KO00040 Pentose and glucuronate interconversions | 84 |
| KO00051 Fructose and mannose metabolism | 138 |
| KO00052 Galactose metabolism | 79 |
| KO00053 Ascorbate and aldarate metabolism | 78 |
| KO00500 Starch and sucrose metabolism | 179 |
| KO00520 Amino sugar and nucleotide sugar metabolism | 256 |
| KO00620 Pyruvate metabolism | 622 |
| KO00630 Glyoxylate and dicarboxylate metabolism | 641 |
| KO00640 Propanoate metabolism | 446 |
| KO00650 Butanoate metabolism | 394 |
| KO00660 C5-Branched dibasic acid metabolism | 141 |
| KO00562 Inositol phosphate metabolism | 93 |
| **Amino acid metabolism** | **3472** |
| KO00250 Alanine, aspartate and glutamate metabolism | 402 |
| KO00260 Glycine, serine and threonine metabolism | 428 |
| KO00270 Cysteine and methionine metabolism | 419 |
| KO00280 Valine, leucine and isoleucine degradation | 415 |
| KO00290 Valine, leucine and isoleucine biosynthesis | 200 |
| KO00300 Lysine biosynthesis | 153 |
| KO00310 Lysine degradation | 228 |
| KO00220 Arginine biosynthesis | 208 |
| KO00330 Arginine and proline metabolism | 254 |
| KO00340 Histidine metabolism | 166 |
| KO00350 Tyrosine metabolism | 148 |
| KO00360 Phenylalanine metabolism | 193 |
| KO00380 Tryptophan metabolism | 246 |
| KO00400 Phenylalanine, tyrosine and tryptophan biosynthesis | 220 |
| **Energy metabolism** | **1602** |
| KO00190 Oxidative phosphorylation | 677 |
| KO00195 Photosynthesis | 151 |
| KO00196 Photosynthesis - antenna proteins | 35 |
| KO00710 Carbon fixation in photosynthetic organisms | 260 |
| KO00910 Nitrogen metabolism | 232 |
| KO00920 Sulfur metabolism | 247 |
| **Nucleotide metabolism** | **1506** |
| KO00230 Purine metabolism | 879 |
| KO00240 Pyrimidine metabolism | 627 |
| **Metabolism of cofactors and vitamins** | **1431** |
| KO00730 Thiamine metabolism | 114 |
| KO00740 Riboflavin metabolism | 61 |
| KO00750 Vitamin B6 metabolism | 77 |
| KO00760 Nicotinate and nicotinamide metabolism | 145 |
| KO00770 Pantothenate and CoA biosynthesis | 191 |
| KO00780 Biotin metabolism | 151 |
| KO00785 Lipoic acid metabolism | 24 |
| KO00790 Folate biosynthesis | 126 |
| KO00670 One carbon pool by folate | 148 |
| KO00860 Porphyrin and chlorophyll metabolism | 308 |
| KO00130 Ubiquinone and other terpenoid-quinone biosynthesis | 86 |
| **Lipid metabolism** | **1100** |
| KO00061 Fatty acid biosynthesis | 250 |
| KO00062 Fatty acid elongation | 5 |
| KO00071 Fatty acid degradation | 286 |
| KO00072 Synthesis and degradation of ketone bodies | 90 |
| KO00073 Cutin, suberine and wax biosynthesis | 1 |
| KO00100 Steroid biosynthesis | 7 |
| KO00561 Glycerolipid metabolism | 105 |
| KO00564 Glycerophospholipid metabolism | 188 |
| KO00565 Ether lipid metabolism | 14 |
| KO00600 Sphingolipid metabolism | 5 |
| KO00590 Arachidonic acid metabolism | 25 |
| KO00591 Linoleic acid metabolism | 10 |
| KO00592 alpha-Linolenic acid metabolism | 20 |
| KO01040 Biosynthesis of unsaturated fatty acids | 94 |
| **Metabolism of other amino acids** | **734** |
| KO00410 beta-Alanine metabolism | 150 |
| KO00430 Taurine and hypotaurine metabolism | 68 |
| KO00440 Phosphonate and phosphinate metabolism | 25 |
| KO00450 Selenocompound metabolism | 155 |
| KO00460 Cyanoamino acid metabolism | 70 |
| KO00472 D-Arginine and D-ornithine metabolism | 4 |
| KO00480 Glutathione metabolism | 262 |
| **Biosynthesis of other secondary metabolites** | **263** |
| KO00940 Phenylpropanoid biosynthesis | 42 |
| KO00950 Isoquinoline alkaloid biosynthesis | 33 |
| KO00960 Tropane, piperidine and pyridine alkaloid biosynthesis | 49 |
| KO00232 Caffeine metabolism | 1 |
| KO00966 Glucosinolate biosynthesis | 23 |
| KO00332 Carbapenem biosynthesis | 15 |
| KO00261 Monobactam biosynthesis | 100 |
| **Metabolism of terpenoids and polyketides** | **221** |
| KO00900 Terpenoid backbone biosynthesis | 180 |
| KO00909 Sesquiterpenoid and triterpenoid biosynthesis | 8 |
| KO00906 Carotenoid biosynthesis | 25 |
| KO00908 Zeatin biosynthesis | 8 |
| **Glycan biosynthesis and metabolism** | **45** |
| KO00510 N-Glycan biosynthesis | 17 |
| KO00514 Other types of O-glycan biosynthesis | 2 |
| KO00531 Glycosaminoglycan degradation | 8 |
| KO00563 Glycosylphosphatidylinositol (GPI)-anchor biosynthesis | 3 |
| KO00603 Glycosphingolipid biosynthesis - globo and isoglobo series | 1 |
| KO00604 Glycosphingolipid biosynthesis - ganglio series | 2 |
| KO00511 Other glycan degradation | 12 |
| **Environmental Information Processing** |  |
| **Membrane transport** | **1062** |
| KO02010 ABC transporters | 1062 |
| **Signal transduction** | **71** |
| KO04075 Plant hormone signal transduction | 16 |
| KO04070 Phosphatidylinositol signaling system | 55 |
| KO04016 MAPK signaling pathway - plant | 80 |
| **Genetic Information Processing** |  |
| **Translation** | **4088** |
| KO03010 Ribosome | 2883 |
| KO00970 Aminoacyl-tRNA biosynthesis | 724 |
| KO03013 RNA transport | 89 |
| KO03015 mRNA surveillance pathway | 35 |
| KO03008 Ribosome biogenesis in eukaryotes | 357 |
| **Folding, sorting and degradation** | **802** |
| KO03060 Protein export | 216 |
| KO04141 Protein processing in endoplasmic reticulum | 114 |
| KO04130 SNARE interactions in vesicular transport | 8 |
| KO04120 Ubiquitin mediated proteolysis | 44 |
| KO04122 Sulfur relay system | 63 |
| KO03050 Proteasome | 33 |
| KO03018 RNA degradation | 324 |
| **Replication and repair** | **801** |
| KO03030 DNA replication | 165 |
| KO03410 Base excision repair | 125 |
| KO03420 Nucleotide excision repair | 162 |
| KO03430 Mismatch repair | 166 |
| KO03440 Homologous recombination | 176 |
| KO03450 Non-homologous end-joining | 7 |
| **Transcription** | **301** |
| KO03020 RNA polymerase | 176 |
| KO03022 Basal transcription factors | 5 |
| KO03040 Spliceosome | 120 |
| **Cellular Processes** |  |
| **Transport and catabolism** | **256** |
| KO04144 Endocytosis | 65 |
| KO04145 Phagosome | 46 |
| KO04146 Peroxisome | 145 |
| KO04136 Autophagy - other | 13 |
| **Organismal Systems** |  |
| **Environmental adaptation** | **62** |
| KO04712 Circadian rhythm - plant | 3 |
| KO04626 Plant-pathogen interaction | 59 |
| **Human Diseases** |  |
| **Drug resistance** | **43** |
| KO01502 Vancomycin resistance | 43 |

### Differential expression analysis between WT and Mutant G1-C1 strain

To have a comprehensive understanding of the molecular mechanism of a higher astaxanthin level in the mutant strain, the differentially expressed transcripts between WT and mutant G1-C1 strains of *Coelastrum* sp. were further evaluated. There was a total of 87,892 DEGs according to the criteria of fold change greater than two and *q*-value less than 0.05. Among these genes, 70,119 genes were upregulated, and 17,773 were downregulated (**Fig 6**). The upregulated unigenes were further classified into various categories using GO functional enrichment analysis and KEGG pathway enrichment analyses. Enrichment analysis was performed to further characterize the functions of DEGs and determine the DEGs involved in the most important biochemical metabolic pathways that are significantly enriched against the transcriptome background [34].

**Fig 6.**
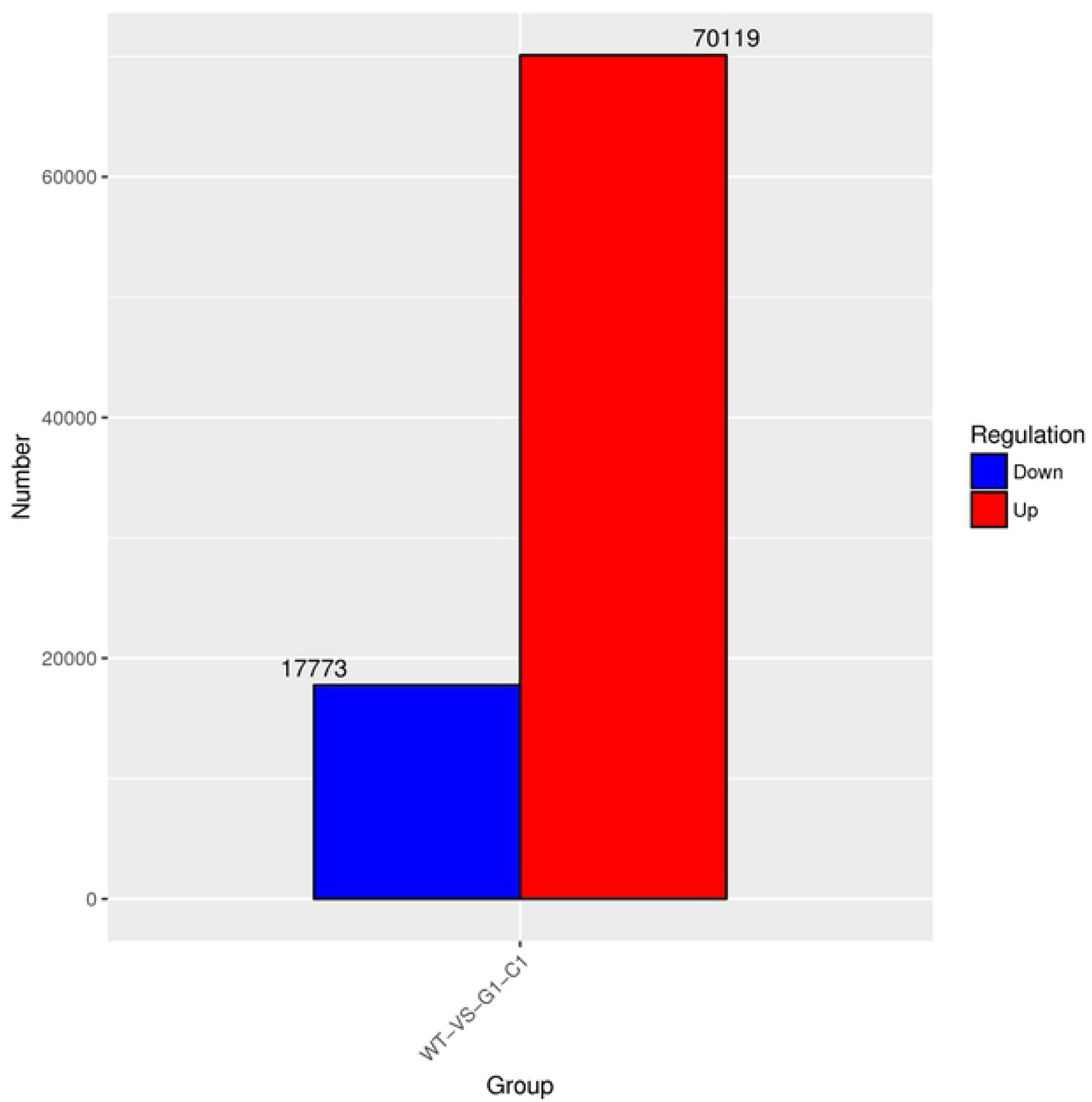
The numbers of DEGs of *Coelastrum* sp. between WT and mutant G1-C1 strains

### GO and KEGG enrichment analysis of DEGs

To identify the functional roles of the identified DEGs in more detail, GO functional enrichment analysis of the DEGs were performed. DEGs were classified into three functional categories, including BP, CC, and MF. As shown in **Fig 7**, GO enrichment analysis revealed that 61 GO terms were significantly enriched in the DEGs responding to mutant G1-C1 of *Coelastrum* sp. Compared with the WT, mutant G1-C1 affected diverse functional categories in MF, BP, and CC categories. The MF group was enriched in 19 subcategories with ‘catalytic activity’, and ‘binding’ which represented the most common categories in upregulated DEGs. The genes in the CC category were enriched into 20 subcategories where the ‘cell part’ represented majority of this category. While predicted proteins assigned to the BP category were assigned to 22 subcategories mainly enriched in the ‘metabolic process’ followed by the ‘cellular process’.

**Fig 7.**
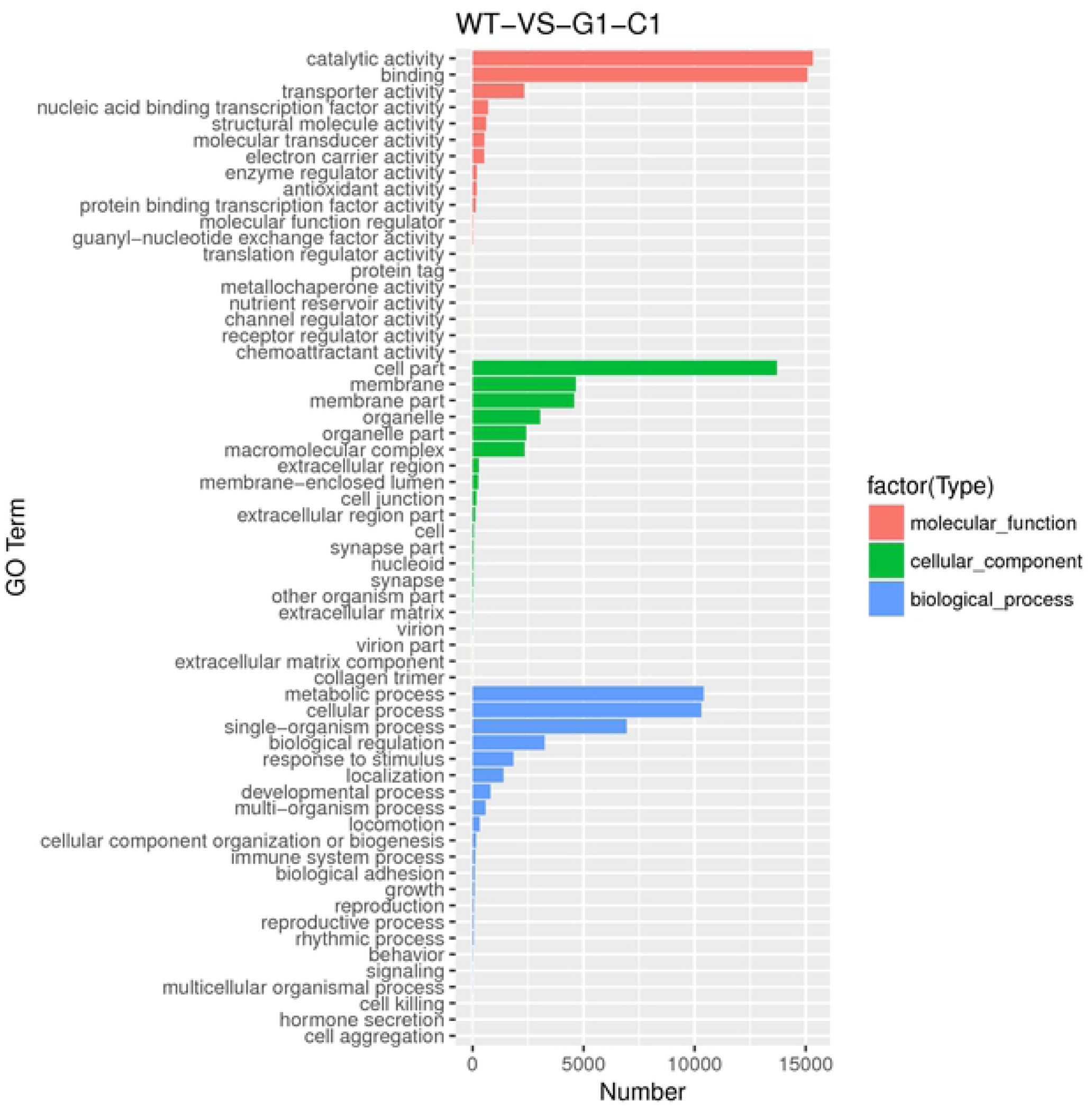
GO enrichment analysis of DEGs. X-axis: Number of DEGs in the GO category. Y-axis: GO term. Color code is to distinguish the categories into the molecular function, cellular component, and biological processes

All these findings showed that a large fraction of upregulated DEGs were enriched in catalytic activity, cell part, metabolic processes, and cellular processes, thus suggesting possible mechanisms which correspond to mutant G1-C1 *Coelastrum* sp. Results have shown that these transcripts could be the genes enriched in various metabolic pathways involved in the cell and enzyme activity to adapt to the cell physiology of mutant G1-C1 *Coelastrum* sp. Besides, the increase in biomass of mutant G1-C1 may also be correlated with the enriched biological process at the cellular level that affects the cellular physiological process and growth of cells.

The KEGG pathway enrichment analysis was used to identify the biological pathways of identified DEGs. This technique was used to determine pathways that are significantly represented in the gene list of interests. The scatter plot in **Fig 8** shows the degree of KEGG enrichment measured by the Rich factor, *q*-value, and the number of genes enriched in this pathway. The scatter plot contains the top 30 pathways that are most significantly enriched. The *q*-value is the corrected *p*-value ranging from 0 to 1. The closer to zero, the more significant enrichment. The *q*-value < 0.5 was set as the threshold for differential expression. A lower value indicates more significant pathway enrichment. As shown in **Fig 8**, transcriptomic analysis of *Coelastrum* sp. mutant showed genome-wide gene expression changes related to different metabolic processes. In particular, a large set of DEGs that were upregulated involved in the mutant G1-C1 strain were mostly enriched in the “Biosynthesis of secondary metabolites” and “Metabolic pathways”. Likely, the genetically modified *Coelastrum* sp. mutant G1-C1 using chemical mutagenesis may possess mutated enzymes with an altered expression. Therefore, these results suggested that mutant G1-C1 may have enhanced multiple metabolic pathways and secondary metabolites involved in *Coelastrum* sp. as compared to the WT. These DEGs involved in the KEGG pathway and GO analysis might probably be the candidate genes involved in carotenoid biosynthesis. Therefore, further studies are required to determine the role of these genes.

**Fig 8.**
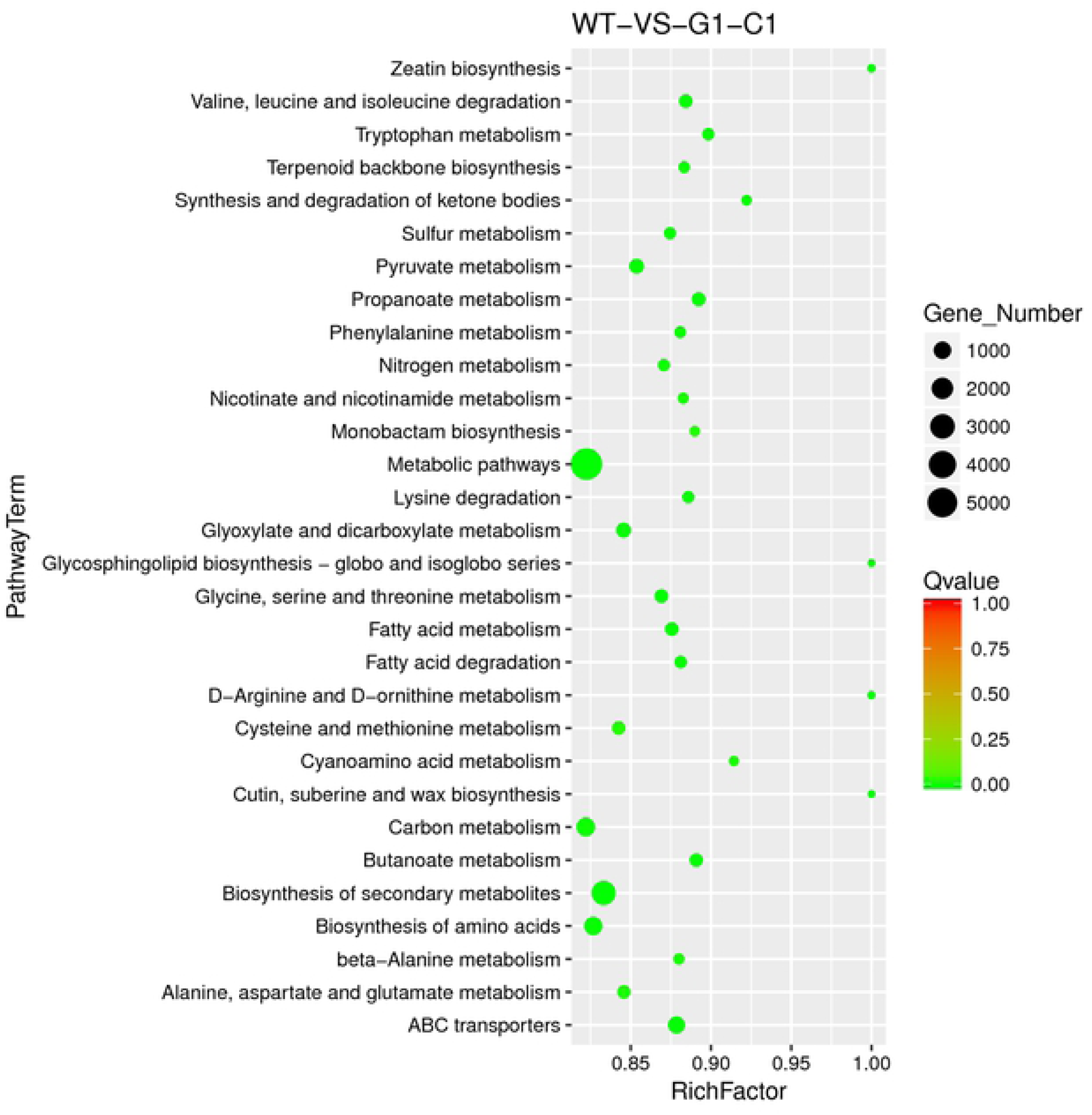
Scatter plot depicting the KEGG enrichment categories of DEGs between mutant G1-C1 and wild type strains. X-axis: Rich Factor. Y-axis: KEGG pathways. The rich factor is the ratio of DEGs enriched in the pathway to all genes. The size of the dots is positively correlated to the number of DEGs in the pathway. Different colors of the points indicate different *q*-value ranges

### Transcriptome and pathway analysis involved in astaxanthin biosynthesis

The accumulation of astaxanthin in the mutant strain obtained by chemical mutagenesis (G1-C1) yielded high astaxanthin levels compared to the WT strain. To explore the molecular mechanisms of higher astaxanthin contents in the mutant, the differences of gene expression in carotenoid biosynthesis between mutant G1-C1 and WT strains were analyzed using RNA-seq. Particularly, differential expressions of genes involved in carotenoid biosynthesis would provide strong evidence of the exact mechanisms responsible for altering astaxanthin production in *Coelastrum* sp. mutant. Therefore, genes of interest that are specifically involved in carotenoid biosynthesis were further investigated, and significant differential expression in the mutant was observed.

In microalgae, the genes involved in carotenoid and astaxanthin biosynthesis are regulated by a series of carotenogenic genes [35, 36]. Transcript levels of genes involved in carotenoid and astaxanthin biosynthesis were determined using transcriptomic analysis. Based on the transcriptomic data shown in **Table 3**, several unigenes were differentially expressed in the mutant strain compared to the WT strain. There are three steps involved in the astaxanthin biosynthesis pathway in microalgae. The first step is to synthesize isopentenyl pyrophosphate (IPP) which is known as β-carotene precursor either via the 2-C-methyl-d-erythritol-4-phosphate (MEP) pathway in the plastid or 3,5-dihydroxy-3-methyl-valonate (MVA) pathway in the cytoplasm. The second step is to synthesize β-carotene and followed by synthesizing astaxanthin as the final step [37–39].

**Table 3.**
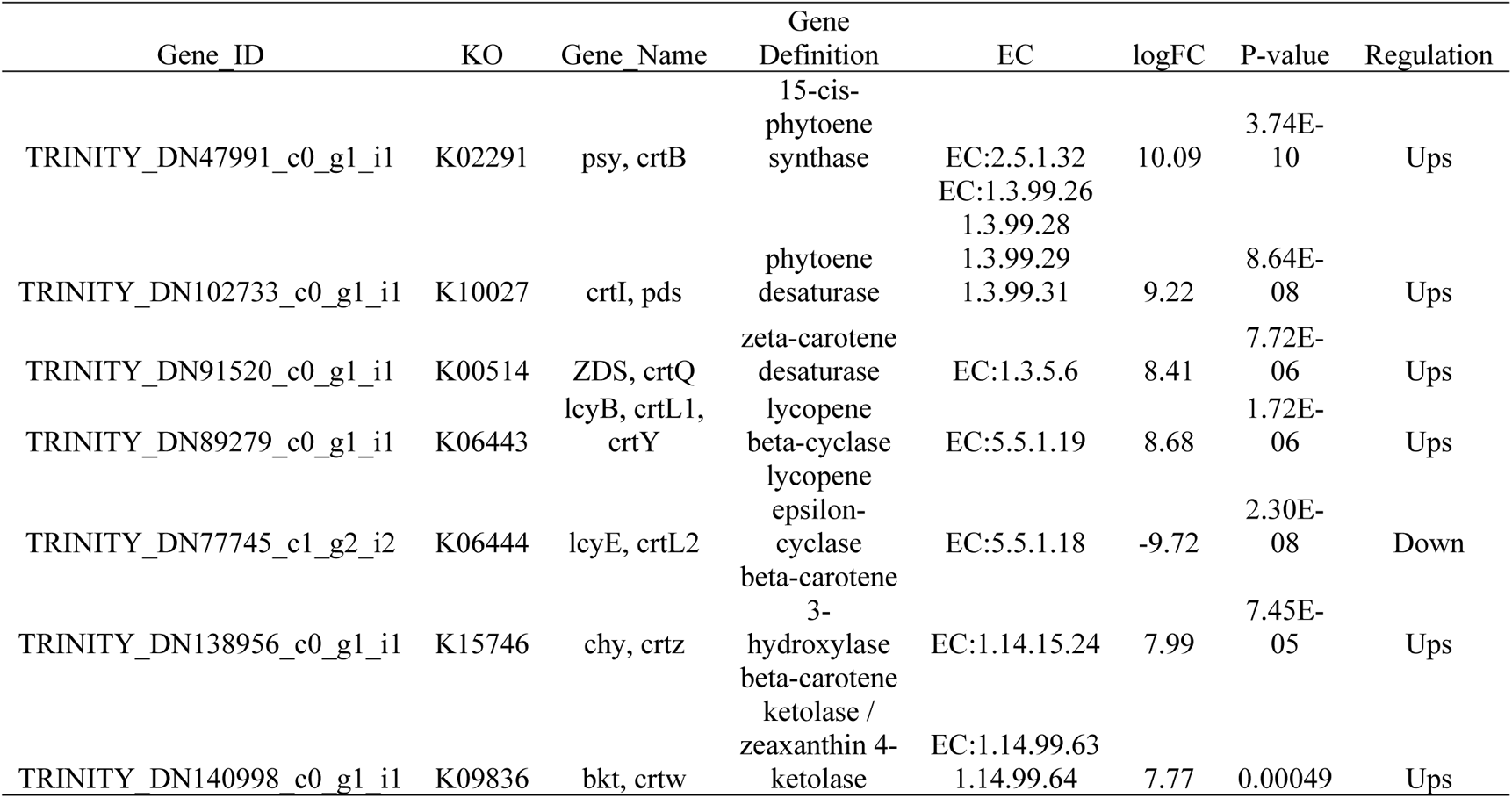
Transcription changes of significantly different unigenes related to astaxanthin biosynthesis pathway of mutant G1-C1 compared to WT of *Coelastrum* sp.

IPP is an important intermediate of the MEP and MVA pathways. In microalgae, IPP can be synthesized either via MEP or MVA pathway. However, based on the transcriptomic data of *Coelastrum* sp., only the genes involved in the MEP pathway were identified. The isomerization of IPP to dimethylallyl pyrophosphate (DMAPP) was catalyzed by IPP isomerase (IPI). It was found that the expression level of *ipi* can affect the metabolic of precursor substances in the carotenoid synthesis pathway and affect the downstream metabolic pathway [40]. The IPP synthesis in the MEP pathway was started with pyruvate and glyceraldehyde-3-phosphate. In this study, eight different enzymes involved in the MEP pathway were identified in the mutant G1-C1 strain (**Fig 9**). According to the transcriptomic analysis of *Coelasrum* sp., the expression of the eight different genes in the MEP pathway were significantly upregulated at different levels, providing IPP for the subsequent astaxanthin synthesis.

**Fig 9.**
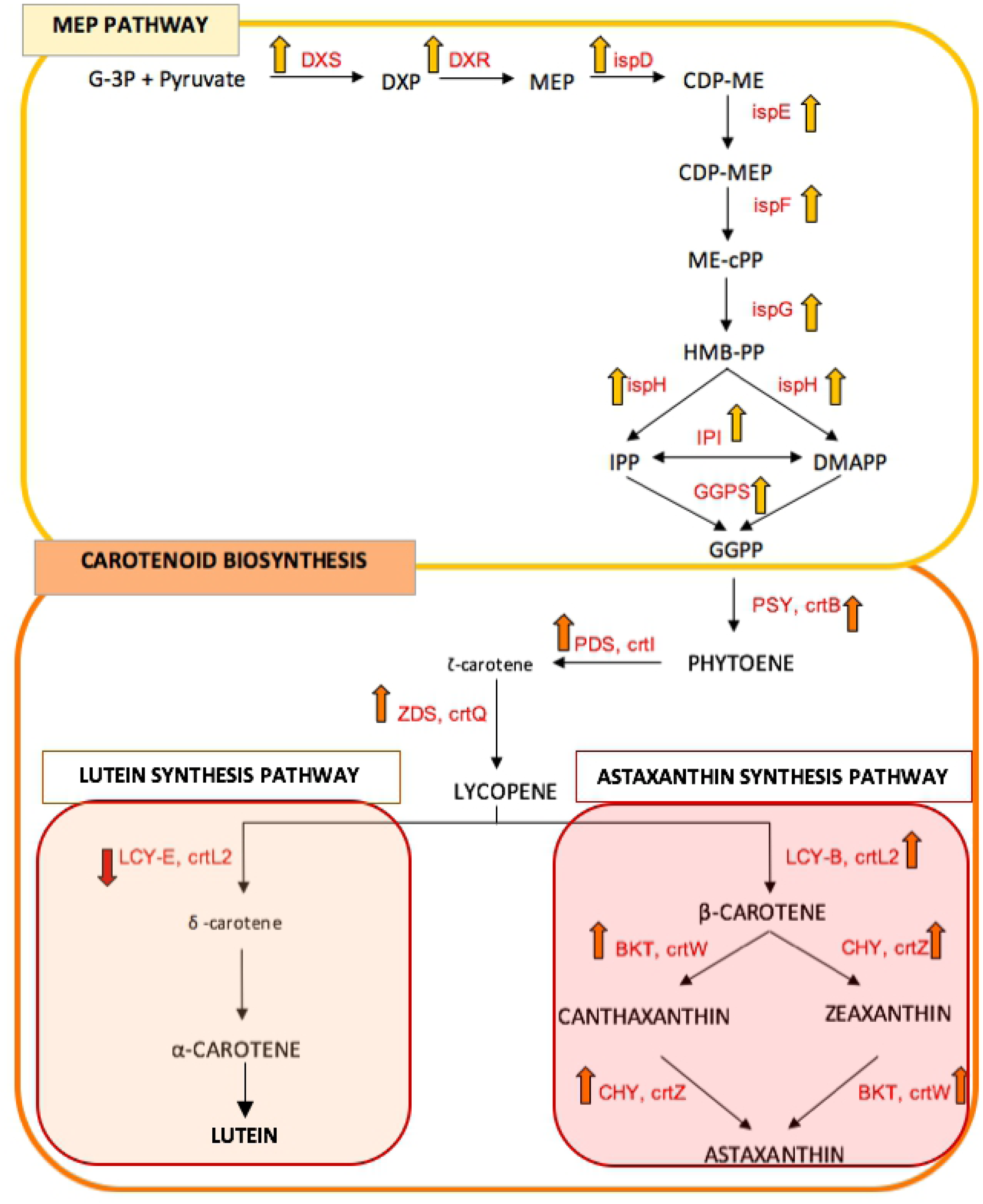
Transcription regulation of genes involved in the astaxanthin biosynthesis pathway of mutant G1-C1 compared to WT *Coelastrum* sp. The upwards and downward solid arrows indicate the upregulation and downregulation of respective genes. DXS: 1-deoxy-D-xylulose-5-phosphatesynthase; DXR: 1-deoxy-D-xylulose-5-phosphate reductoisomerase; ispD: 2-C-methyl-D-erythritol 4-phosphate cytidylyltransferase; ispE: 4-diphosphocytidyl-2-C-methyl-D-erythritol kinase; ispF: 2-C-methyl-D-erythritol 2,4-cyclodiphosphate synthase; ispG: 4-hydroxy-3-methylbut-2-enyl-diphosphate synthase; ispH: 4-hydroxy-3-methylbut-2-en-1-yl diphosphate reductase; IPI: Isopentenyl pyrophosphate isomerase; GGPS: geranylgeranyl diphosphate synthase; PSY: Phytoene synthase; PDS: phytoene desaturase; ZDS: zeta-carotene desaturase; LCY-B: lycopene beta-cyclase; LCY-E: lycopene epsilon-cyclase; CHY: beta carotene hydroxylase; BKT: beta carotene ketolase

β-carotene is the precursor of astaxanthin synthesis, which was catalyzed in the next step of astaxanthin synthesis. The enzymes involved in the conversion of geranylgeranyl pyrophosphate (GGPP) into β-carotene was successively catalyzed by phytoene synthase (PSY), phytoene desaturase (PDS), and *ζ*-carotene desaturase (ZDS). In this study, the transcriptomic analysis showed that the gene expressions of these genes were upregulated at significantly different levels. Compared with the WT, the upregulations of *psy*, *pds*, and *zds* reported in the mutant were upregulated by 10-, 9.2-, and 8.4-fold, respectively (**Table 3**).

Lycopene is the intermediate substrate for astaxanthin and lutein synthesis. It can be catalyzed by Lycopene β-cyclase (LCY-b) into β-carotene which is a crucial enzyme in the astaxanthin synthesis pathway. Lycopene could also be catalyzed by lycopene ε-cyclase (LCY-e) to generate α-carotene in the lutein synthesis pathway. However, the lutein synthesis pathway can compete with the astaxanthin synthesis pathway for LCY-b and thus reduce astaxanthin synthesis. According to the transcriptomic analysis of *Coelastrum* sp. the upregulation of LCY-b gene expression was accompanied by the downregulation of LCY-e gene expression. LCY-e gene was downregulated by 9.7-fold in mutant G1-C1, while the LCY-b gene was upregulated by 8.7-fold (**Fig 9**). This situation would decrease the competition in lutein metabolism for the lycopene, implying more carbon flow to astaxanthin synthesis rather than lutein. Therefore, the inhibition of a competitive pigment synthesis pathway was probably a reason why mutant G1-C1 of *Coelastrum* sp. could enrich astaxanthin in large quantities as compared to the WT. The upregulation of *lcy-b* accompanied by the downregulation of *lcy-e* was also reported in the study of transcriptome analysis [41, 42].

There are two pathways involved in astaxanthin synthesis from β-carotene; oxidation and hydroxylation of β-carotene [43]. In the last step of astaxanthin synthesis, the production of astaxanthin via canthaxanthin and zeaxanthin could be catalyzed by beta-carotene 3-hydroxylase (CHY/crtz) and beta-carotene ketolase (BKT/crtw). These enzymes have been reported to play a vital role in astaxanthin biosynthesis by mediating the final enzymatic conversions leading to astaxanthin synthesis [44, 45]. In particular, the transcriptomic analysis of *Coelastrum* sp. showed that the transcription level of *bkt* and *chy* genes that encoded for the rate-limiting steps of astaxanthin were both upregulated in mutant G1-C1 by 7.8- and 8.0-fold, respectively. The results show that the upregulation of *bkt* and *chy* expression levels in the mutant G1-C1 correlated to the higher astaxanthin content in the mutant compared to the WT.

## Conclusions

Mutant obtained through chemical mutagenesis (Mutant G1-C1) was observed to enhance astaxanthin accumulation in a newly isolated *Coelastrum* sp. However, its molecular mechanisms are yet to be identified. Comparative transcriptome analysis provides new insight into *Coelastrum* sp. with molecular mechanisms of astaxanthin biosynthesis underlying the mutant G1-C1 and WT. A total of 87,892 DEGs were identified where 70,119 genes were upregulated, and 17,773 genes were downregulated. Functional analysis indicated that multiple genes in different functional categories were differentially expressed. Notably, the KEGG analysis implied that the changes in the metabolic pathway and biosynthesis of secondary metabolites were most prominent. Using transcriptome sequencing, this study analyzed the expression patterns of genes involved in carotenoid and astaxanthin biosynthesis. Astaxanthin biosynthesis in *Coelastrum* sp. mutant (G1-C1) was accelerated by upregulation of genes involved in β-carotene synthesis (PSY, PDS, ZDS, and LCY-b) and astaxanthin synthesis (BKT and CHY). Besides, the downregulation of the LCY-e gene has suppressed the lutein pathway biosynthesis and thus, promoted astaxanthin synthesis. In summary, these genes play a prominent role in astaxanthin biosynthesis and could result in enhanced astaxanthin accumulation in the mutant G1-C1 of *Coelastrum* sp.

## Author Contributions

AT carried out the experimental work, analysed the data and prepared the manuscript. All authors contributed intellectually to the presented work and approved the final version of the manuscript.

## Acknowledgments

The authors are grateful to the Malaysia-Japan International Institute of Technology (MJIIT), University of Tsukuba, Japan, and Japan Student Services Organization (JASSO) scholarship for providing financial supports and technical facilities for the study to be carried out.

## References

1. Dragos N, Bercea V, Bica A, Druga B, Nicoara A, Coman C. Astaxanthin production from a new strain of *Haematococcus pluvialis* grown in batch culture. Ann RSCB. 2016; 2: 353–361.

2. Shah MMR, Liang Y, Cheng JJ, Daroch M. Astaxanthin-producing green microalga *Haematococcus pluvialis*: from single cell to high value commercial products. Front Plant Sci. 2016; 7: 1–28. https://doi.org/10.3389/fpls.2016.00531

3. Fakhri S, Abbaszadeh F, Dargahi L, Jorjani M. Astaxanthin: A mechanistic review on its biological activities and health benefits. Pharmacol Res. 2018; 136: 1–20. https://doi.org/10.1016/j.phrs.2018.08.012

4. Ng QX, De Deyn MZQ, Loke W, Foo NX, Chan HW, Yeo WS. Effects of astaxanthin supplementation on skin health: a systematic review of clinical studies. J Diet Suppl. 2020; 1–14. https://doi.org/10.1080/19390211.2020.1739187

5. Nouchi R, Suiko T, Kimura E, Takenaka H, Murakoshi M, Uchiyama A, et al. Effects of lutein and astaxanthin intake on the improvement of cognitive functions among healthy adults: a systematic review of randomized controlled trials. Nutrients. 2020; 12(3): 617. https://doi.org/10.3390/nu12030617

6. Orosa M, Franqueira D, Cid A., Abalde J. Carotenoid accumulation in *Haematococcus pluvialis* in mixotrophic growth. Biotechnol Lett (2001); 23: 373–378. https://doi.org/10.1023/A:1005624005229

7. Liu J, Sun Z, Gerken H, Liu Z, Jiang Y, Chen F. *Chlorella zofingiensis* as an alternative microalgal producer of astaxanthin: biology and industrial potential. Mar Drugs. 2014; 12: 3487–3515. https://doi.org/10.3390/md12063487

8. Tharek A, Jamaluddin H, Salleh MM, Yahya NA, Kaha M, Hara H, et al. Astaxanthin production by tropical microalgae strains isolated from environment in Malaysia. Asian J Microbiol Biotechnol Environ Sci. 2020a; 22(1): 168–173. https://doi.org/10.6084/m9.figshare.12972881.v1

9. Tharek A, Mohamad SE, Iwamoto K, Suzuki I, Hara H, Dolah R, et al. Enhanced astaxanthin production by oxidative stress using methyl viologen as a reactive oxygen species (ROS) reagent in green microalgae *Coelastrum* sp. Indones J Biotechnol. 2020b; 25(2): 95–101. https://doi.org/10.22146/ijbiotech.54092

10. Tharek A, Yahya A, Salleh MM, Jamaluddin H, Yoshizaki S, Dolah R, et al. Improvement of astaxanthin production in *Coelastrum* sp. by optimization using Taguchi Method. Appl Food Biotechnol. 2020c; 7(4): 205–214. https://doi.org/10.22037/afb.v7i4.29697

11. Kilian O, Benemann CS, Niyogi KK, Vick B. High-efficiency homologous recombination in the oil-producing alga *Nannochloropsis* sp. Proc Natl Acad Sci. USA. 2011; 108(52): 21265–21269. https://doi.org/10.1073/pnas.1105861108

12. Chaturvedi R, Uppalapati S, Alamsjah M, Fujita Y. Isolation of quizalofop-resistant mutants of *Nannochloropsis oculata* (Eustigmatophyceae) with high eicosapentaenoic acid following N-methyl-N-nitrosourea-induced random mutagenesis. J Appl Phycol. 2004; 16: 135–144. https://doi.org/10.1023/B:JAPH.0000044826.70360.8e

13. Sandesh Kamath B, Vidhyavathi R, Sarada R, Ravishankar GA. Enhancement of carotenoids by mutation and stress induced carotenogenic genes in *Haematococcus pluvialis* mutants. Bioresour Technol. 2008; 99(18): 8667–8673. https://doi.org/10.1016/j.biortech.2008.04.013

14. Hong ME, Choi SP, Park Y, Kim Y, Chang WS, Kim BW, et al. Astaxanthin production by a highly photosensitive *Haematococcus* mutant. Process Biochem. 2012; 47: 1972– 1979. https://doi.org/10.1016/j.procbio.2012.07.007

15. Sharon-Gojman R, Maimon E, Leu S, Zarka A, Boussiba S. Advanced methods for genetic engineering of *Haematococcus pluvialis* (Chlorophyceae, Volvocales). Algal Res. 2015; 10: 8–15. https://doi.org/10.1016/j.algal.2015.03.022

16. Fischer R. Isolation of mutants, a key for the analysis of complex pathways and for strain improvement. In: Verma A, editor. Microbes for Health, Wealth and Sustainable Environment. Malhotra Publishing House, New Delhi, India; 1998. pp. 739–751.

17. Chen Y, Li D, Lu W, Xing J, Hui B, Han Y. Screening and characterization of astaxanthin-hyperproducing mutants of *Haematococcus pluvialis*. Biotechnol Lett. 2003; 25: 527–529. https://doi.org/10.1023/A:1022877703008

18. Gómez PI, Inostroza I, Pizarro M, Pérez J. From genetic improvement to commercial-scale mass culture of a Chilean strain of the green microalga *Haematococcus pluvialis* with enhanced productivity of the red ketocarotenoid astaxanthin. AoB PLANTS. 2013; 5: 1–7. https://doi.org/10.1093/aobpla/plt026

19. Liu Z, Liu C, Hou Y, Chen S, Xiao D, Zhang J, et al. Isolation and characterization of a marine microalgae for biofuel production with astaxanthin as a co-product. Energies. 2013; 6(6): 2759–2772. https://doi.org/10.3390/en6062759

20. Ubeda B, Galvez JA, Michel M, Bartual A. Microalgae cultivation in urban wastewater: *Coelastrum cf. pseudomicroporum* as a novel carotenoid source and a potential microalgae harvesting tool. Bioresour Technol. 2017; 228: 210–217. https://doi.org/10.1016/j.biortech.2016.12.095

21. Gwak Y, Hwang YS, Wang B, Kim M, Jeong J, Lee CG, et al. Comparative analyses of lipidomes and transcriptomes reveal a concerted action of multiple defensive systems against photooxidative stress in *Haematococcus pluvialis*. J exp Bot. 2014; 65(15): 4317–4334. https://doi.org/10.1093/jxb/eru206

22. Wang L, Feng Z, Wang X, Zhang X. DEGseq: An R package for identifying differentially expressed genes from RNA-seq data. Bioinform. 2010; 26: 136–138. https://doi.org/10.1093/bioinformatics/btp612

23. Kato S. Laboratory culture and morphology of *Colacium vesiculosum* Ehrb. (Euglenophyceae). Jpn J Phycol. 1982; 30: 63–67.

24. Boussiba S, Vonshak A. Astaxanthin accumulation in the green alga *Haematococcus pluvialis*. Plant Cell Physiol. 1991; 32: 1077–1082. https://doi.org/10.1093/oxfordjournals.pcp.a078171

25. Lichtenthaler HK. Chlorophylls and carotenoids: pigments of photosynthetic biomembranes, In: Packer L, Douce R, editors. Methods Enzymol; 1987. 148: pp. 350–382. https://doi.org/10.1016/0076-6879(87)48036-1

26. Martin M. Cutadapt removes adapter sequences from high-throughput sequencing reads. EMBnet J. 2011; 17(1): 10–12. https://doi.org/10.14806/ej.17.1.200

27. Grabherr MG, Haas BJ, Yassour M, Levin JZ, Thompson DA, Amit I, et al. Full-length transcriptome assembly from RNA-seq data without a reference genome. Nat Biotechnol. 2011; 29(7): 644–52. https://doi.org/10.1038/nbt.1883

28. Langmead B, Trapnell C, Pop M, Salzberg SL. Ultrafast and memory-efficient alignment of short DNA sequences to the human genome. Genome Biol. 2009; 10: R25. https://doi.org/10.1186/gb-2009-10-3-r25

29. Fu L, Niu B, Zhu Z, Wu S, Li W. CD-HIT: accelerated for clustering the next-generation sequencing data. Bioinform. 2012; 28(23): 3150–31522. https://doi.org/10.1093/bioinformatics/bts565

30. Li B, Dewey CN. RSEM: accurate transcript quantification from RNA-Seq data with or without a reference genome. BMC Bioinform. 2011; 12: 323. https://doi.org/10.1186/1471-2105-12-323

31. Anders S, Huber W. Differential expression of RNA-Seq data at the gene level-the DESeq package (DESeq). 2012.

32. Kanehisa M, Goto S, Furumichi M, Tanabe M, Hirakawa M. KEGG for representation and analysis of molecular networks involving diseases and drugs. Nucleic Acids Res. 2010; 38: 355–360. https://doi.org/10.1093/nar/gkp896

33. Huang Y, Xiong JL, Gao X, Sun X. Transcriptome analysis of the Chinese giant salamander (*Andrias davidianus*) using RNA-sequencing. Genom. Data. 2017; 14: 126–131. https://doi.org/10.1016/j.gdata.2017.10.005

34. Kanehisa M, Araki M, Goto S, Hattori M, Hirakawa M, Itoh M. KEGG for linking genomes to life and the environment. Nucleic Acids Res. 2007; 36: 480–484. https://doi.org/10.1093/nar/gkm882

35. Lee C, Choi YE, Yun YS. A strategy for promoting astaxanthin accumulation in *Haematococcus pluvialis* by 1-aminocyclopropane-1-carboxylic acid application. J Biotechnol. 2016; 236: 120–127. https://doi.org/10.1016/j.jbiotec.2016.08.012

36. Wen Z, Liu Z, Hou Y, Liu C, Gao F, Zheng Y, et al. Ethanol induced astaxanthin accumulation and transcriptional expression of carotenogenic genes in *Haematococcus pluvialis*. Enzyme Microb Technol. 2015; 78: 10–17. https://doi.org/10.1016/j.enzmictec.2015.06.010

37. Vidhyavathi R, Venkatachalam L, Sarada R, Ravishankar GA. Regulation of carotenoid biosynthetic genes expression and carotenoid accumulation in the green alga *Haematococcus pluvialis* under nutrient stress conditions. J Exp Bot. 2008; 59(6): 1409–1418. https://doi.org/10.1093/jxb/ern048

38. Stange C. Carotenoids in nature. vol. 8. Basel: Springer International Publishing; 2016. pp. 219–231. https://doi.org/10.1007/978-3-319-39126-7

39. Vranova E, Coman D, Gruissem W. Network analysis of the MVA and MEP pathways for isoprenoid synthesis. Annu Rev Plant Biol. 2013; 64: 665–700. https://doi.org/10.1146/annurev-arplant-050312-120116

40. Chappell J. The biochemistry and molecular biology of isoprenoid metabolism. Plant Physiol. 1995; 107(1): 1–6. https://doi.org/10.1104/pp.107.1.1

41. He B, Hou L, Dong M, Shi J, Huang X, Ding Y, et al. Transcriptome analysis in *Haematococcus pluvialis*: astaxanthin induction by high light with acetate and Fe^2+^. Int. J Mol Sci. 2018; 19: 175–193. https://doi.org/10.3390/ijms19010175

42. Gao Z, Miao X, Zhang X, Wu G, Guo Y, Wang M, et al. Comparative fatty acid transcriptomic test and iTRAQ-based proteomic analysis in *Haematococcus pluvialis* upon salicylic acid (SA) and jasmonic acid (JA) inductions. Algal Res. 2016; 17: 277– 284. https://doi.org/10.1016/j.algal.2016.05.012

43. Kobayashi M. Astaxanthin biosynthesis enhanced by reactive oxygen species in the green alga *Haematococcus pluvialis*. Biotechnol Bioprocess Eng. 2003; 8: 322–330. https://doi.org/10.1007/BF02949275

44. Steinbrenner J, Linden H. Light induction of carotenoid biosynthesis genes in the green alga *Haematococcus pluvialis*: regulation by photosynthetic redox control. Plant Mol Biol. 2003; 52: 343–356. https://doi.org/10.1023/A:1023948929665

45. Zhong YJ, Huang JC, Liu J, Li Y, Jiang Y, Xu ZF, et al. Functional characterization of various algal carotenoid ketolases reveals that ketolating zeaxanthin efficiently is essential for high production of astaxanthin in transgenic Arabidopsis. J Exp Bot. 2011; 62: 3659–3669. https://doi.org/10.1093/jxb/err070

